# Evolutionary mismatch between nuclear and mitochondrial genomes does not promote reversion mutations in mtDNA

**DOI:** 10.1101/2024.08.21.609033

**Authors:** Melissa Franco, Konstantin Popadin, Dori Woods, Konstantin Khrapko

## Abstract

Serrano et al. (Serrano et al., 2024) use a high-fidelity somatic mtDNA mutation analysis in conplastic mice in which mtDNA was replaced with exogenous mtDNA of different mouse strains. Serrano reported apparent abundant somatic reversion mutations in the exogenous mtDNA that seemed to restore the original mito-nuclear match. If real, such a phenomenon would have important implications for health and genetics. In today’s highly mixed human population, the pairing of potentially mismatched nuclear and mitochondrial genomes is widespread, so the proposed reversion mutagenesis should be commonplace. We demonstrate, however, that these *reversion mutations* are not real but originate from cross-contamination between samples and from NUMTs, the mtDNA pseudogenes located in the nuclear genome.

## The motivating inconsistency and the working hypotheses

Serrano et al. (Serrano et al., 2024) report an apparent extensive enrichment in somatic ‘reversion mutations,’ which allegedly cause exogenous mtDNA from a different mouse strain that has replaced the original mtDNA of B6 mice to ‘evolve’ towards the sequence of the original B6 mtDNA. The ‘enrichment’ of reverse mutations supposedly reflected selective pressure on mtDNA to ‘match’ nuclear DNA. This proposed selection is impossible, however, because most of the ‘reversion mutations’ are synonymous. This inconsistency prompted us to look for alternative hypotheses. Previously (see, e.g., (Khrapko et al., 1994), (Hirano et al., 1997), (Fleischmann et al., 2024)), presumed somatic mutations in mtDNA were often shown to be artifacts derived from NUMTs, i.e., nuclear pseudogenes of mtDNA. Alternatively, false somatic mutations can originate from minor contamination with mtDNA of a different haplotype. Interestingly, both NUMTs and contamination hypotheses predict high synonymity false mutations. High synonymity is indeed a prominent property of ‘reverse mutations’, which contrasts with marked low synonymity of true somatic mutations. We, therefore, evaluated both hypotheses.

### The NUMT hypothesis

NUMTs are mtDNA sequences that have been inserted into nuclear DNA in the course of evolution. NUMTs range from very recent, inserted just a few generations prior, to ancient, dating up to tens of millions of years ago (Popadin et al., 2022). Upon insertion into nDNA, NUMT’s and mtDNA start accumulating different mutations, and their evolutionary paths diverge (**Figure 1**). For a reverse mutation study, it is important that NUMTs preserve the ancestral states for nucleotide positions homologous to those where mutations occurred in mtDNA. In next-generation sequencing, where reads are short (∼130 bp), NUMT-derived reads that overlap ancestral nucleotide positions may be otherwise indistinguishable from mtDNA-derived reads because they do not include any NUMT-specific SNVs, which would identify such reads. In such a case, ancestral nucleotides at polymorphic positions will falsely appear as reverse mutations. This is particularly true for recent pseudogenes (e.g., NUMTchr1, **Figure 1**). Serrano et al. corrected mutant fractions of the ‘reverse mutations’ originating from NUMTchr1 using NUMTchr1-nDNA junctions as a marker. However, other pseudogenes (e.g. NUMTchr2, **Figure 1**) may also contain 130bp-regions fully identical to the B6 mtDNA, so some false mutations may still not be accounted for. Furthermore, most recent NUMTs can potentially be private (as demonstrated in humans (Wei et al., 2020)), i.e., present only in some mouse lineages (e.g. those used by Serrano), and not in the reference mouse genome. The NA are unknown, so their contribution to ‘reverse mutations’ are unknown, so their contribution to ‘reverse mutations’ cannot be quantified.

**Figure 1.**
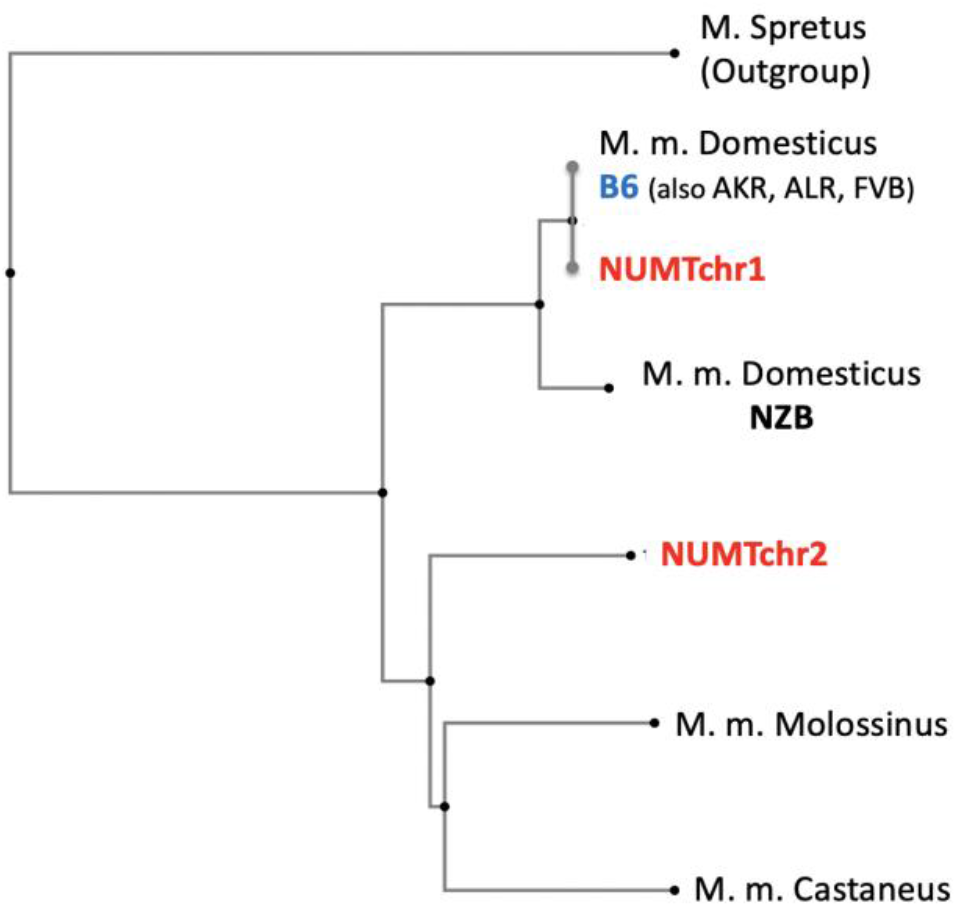
The mouse phylogenetic tree that includes the two NUMTs that may potentially affect the Serrano study. NUMTchr1 on chromosome 1, a fragment of B6 mtDNA was inserted very recently into the B6 nuclear DNA, so there is only one SNV distinguishing NUMT from mtDNA and it’s branch is almost identical to the *B6* branch. NUMTchr2 on chromosome 2, was created ∼1 My ago, the time of separation of mouse subspecies, *Domesticus, Molossinus*, and *Castaneus*.

### The cross-contamination hypothesis

If a mtDNA sample of a certain (‘major’) haplotype is contaminated with mtDNA of a different (‘minor‘) haplotype, sequencing reads derived from the minor haplotype will be perceived as carrying reverse mutations at all SNV positions that belong to the ‘major’ haplotype. Such cross-contamination is indeed potentially possible. Serrano et al. reported one sample with about 25% contamination. 25% is easily detected; however, because reverse mutations in this study were reported at about ∼0.1%, a mere 0.1% contamination would be sufficient to account for them. Such a low contamination would be much more likely to happen and would most likely go undetected.

### The observation of multiple *mutations* per sequencing read supports both the cross-contamination and the NUMT hypotheses

Both above hypotheses make the same specific *testable prediction, especially* for the B6 mouse carrying NZB mtDNA. NZB mtDNA differs by about 90 SNVs from the mtDNA of any of the four other strains used in the study (B6, ALR, AKR, and FVB). Importantly, there are a few ‘SNV-clusters’, where multiple NZB-specific SNVs fall within the length of the Illumina sequencing read (133 bp). Cross-contamination with any non-NZB mtDNA or inclusion of any NUMT sequences covering an SNV-cluster would result in sequencing reads where *all* the multiple NZB-specific SNVs of the cluster appear ‘reversed’ to the B6-specific nucleotides. In contrast, real somatic reverse mutations are rare and should occur independently, leading to nearly zero chances of seeing several of them on the same molecule. To test this prediction, we focused on one area of dense NZB-specific SNV cluster, i.e., 2757-2847 (cluster of 5 SNPs within 90bp). This region does not overlap with NUMTchr1, so any presumed reverse mutations in this region were counted without correction. We obtained sequence reads covering this area from Serrano’s raw data on a typical sample, NZB young brain sample 1, which reportedly contains reverse mutations at about 0.1%. To visualize these ‘reverse mutations’, we used BLAST to map these reads onto the B6 mtDNA query sequence following the approach we used previously (Fleischmann et al., 2024); see Methods for details. BLAST ranges sequences by similarity to the B6 query, so any NZB reads with ‘reverse mutations’ (i.e., mutations towards B6) must conveniently appear at the top of the alignment.

The results of this analysis are presented in **Figure 2**. Consistent with both the contamination and the NUMT hypotheses, the reads containing ‘reverse mutations’ were represented *exclusively* by the fully B6-like reads, i.e., those where *all* 5 SNVs in the cluster were of the B6 haplotype. There are 7 fully B6-like reads and 7,916 NZB reads, i.e., mutant fraction is ∼0.9×10^−3^, in excellent agreement with the Serrano’s estimate for this sample. As argued above, these B6-like reads can be derived either from cross-contamination with a B6 haplotype sample (i.e., B6, ALR, AKR, and FVB), or from a B6-like NUMT, so we conclude that these are NOT ‘reverse mutations’. Moreover, **Figure 2** shows that, remarkably, there are *no* reads with *one* reverse mutation, as expected for somatic reverse mutations. A similar result was obtained for another SNV cluster, i.e. 3 NZB-specific SNVs 4701-81. Again, only fully NZB and fully B6 reads were observed. We conclude that there are no measurable true reverse mutations in the two sequence regions in this sample.

**Figure 2.**
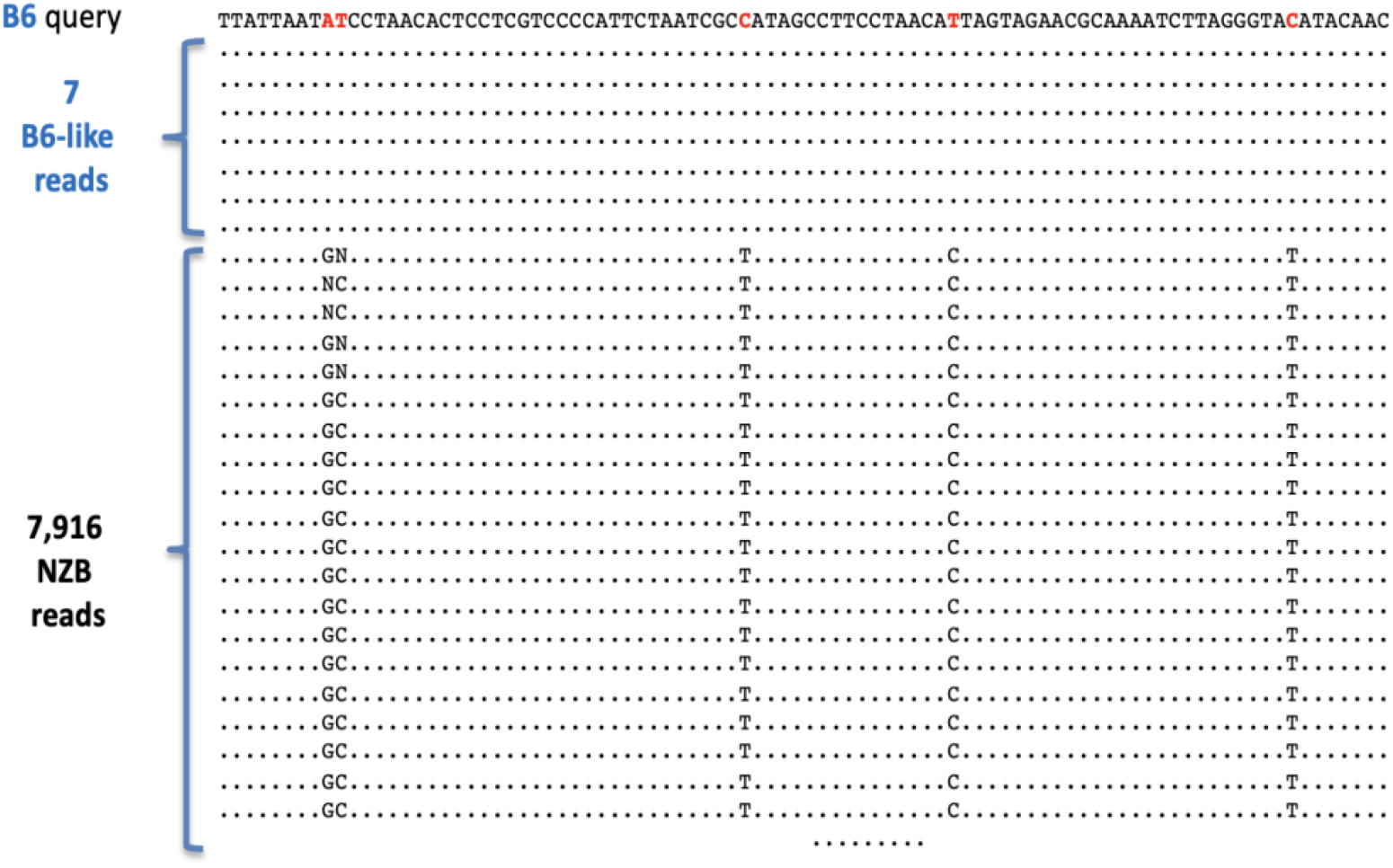
∼8,000 Illumina reads of a sample from NZB/B6 conplastic mouse covering the 2757-2847 region which contains 5 NZB-specific SNVs which are potential sites of the ‘reverse’ NZB>B6 mutations. mtDNA reads were aligned and ranged by extent of identity to the B6 mtDNA query sequence using BLAST (dots depict identity).

### Cross-contamination is the main source of artificial reverse mutations

To determine what is the exact source of false mutations, i.e., NUMT or cross-contamination, we consider that the 2757-2847 region is not overlayed by any known pseudogene in the reference mouse genome (Kent, 2002). This supports cross-contamination as the source. However, it is still possible that ‘reverse mutations’ come from other, private NUMT(s) that are specific to mice used in the Serrao study. We note, however, that NUMT-derived mutations should make a local NUMT ‘footprint’ on mtDNA, whereas cross-contamination of a mouse with NZB mtDNA with any non-NZB mtDNA should result in a global background of ∼90 ‘reverse mutations’ distributed across the entire mitochondrial genome. We, therefore, plotted mutant fractions of reverse mutations along mtDNA of one of the NZB conplastic mouse samples (NZN_Br_Y2) and, indeed, observed a clear footprint covering section of mitochondrial genome homologous to the NUMTchr1 (**Figure 3A**). Importantly, the lack of any other footprint suggests that no other NUMT significantly contributed reverse mutations to this sample (and other samples of this mouse line, which are genetically identical and thus contain the same set of NUMTs, including the private NUMTs!). Likewise, there should be no significant non-NZB contamination, in this particular sample, otherwise there should have been mutational background across the entire mtDNA. NZB_Br_Y2 sample, however, is an unusual non-contaminated sample. A more typical sample, NZB_Br_Y1 (**Figure 3B**), shows a strong uniform background of ‘reverse mutations’ superimposed on the NUMTchr1 footprint. We conclude that sample Y1 is contaminated with B6-like mtDNA. The latter can be from any of the three non-NZB lines used in the study.

**Figure 3.**
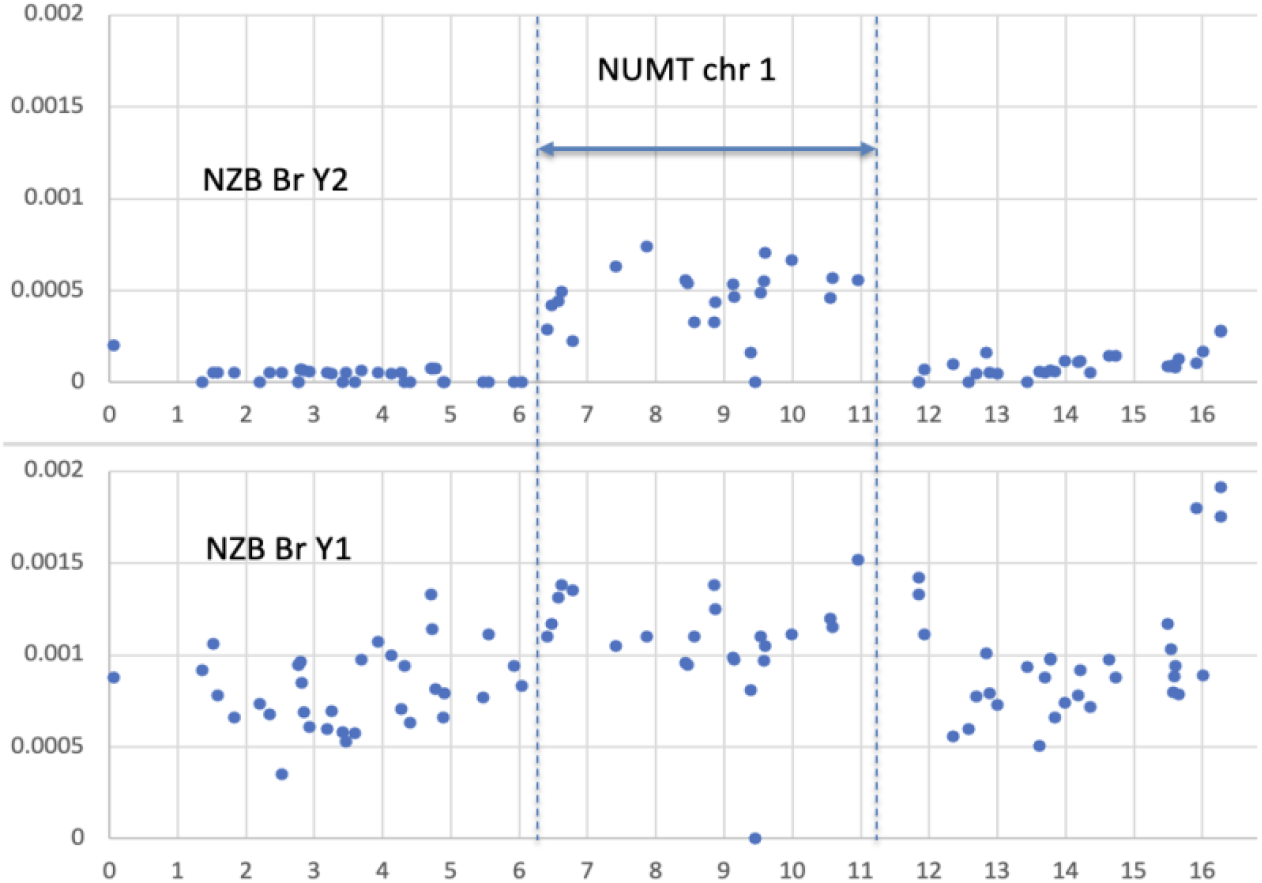
Distribution of ‘reverse mutations’ along the mitochondrial genome in NZB/B6 conplastic mice. **A**. The ‘mutational footprint’ of the NUMTchr1 is clearly visible in an unusual non-contaminated sample. **B**. Cross-contamination with non-NZB mtDNA creates a background of ‘reverse mutations’ overlaying the NUMTchr1 footprint in a more common contaminated sample.

### Concluding remarks

Our study was not meant to exhaustively examine Serrano’s results, which is the prerogative of the authors. Moreover, many of the mutations reported by the authors cannot be tested because of the lack of appropriate markers. Thus, our approach was to show that arbitrarily chosen testable reverse mutations turn out to be artifacts stemming from NUMT and cross-contamination between samples and there is no evidence for the enrichment of somatic reverse mutations.

## Methods

### Data for haplotype analysis (Figure2)

Sequencing reads from the Serrano et al paper were accessed on the Zenodo repository (https://zenodo.org/records/10403641). For each sample analyzed, either the single or double stranded read 1 (read1_sscs or read1_dcs) was used. Though Serrano typically uses double stranded reads in their reported analysis, when comparing the single and double stranded reads in the same loci for the same sample, it became evident that the single stranded reads provided greater depth and therefore provide a better picture of how prevalent the contamination is. The use of single stranded reads for our purposes has no downside. In the double stranded sequencing approach used by Serrano et al., the advantage of using ds reads is that ds reads effectively filter out any single stranded base changes which are not real mutations, but premutagenic damage or instrumental errors of various types. In our case there is no doubt that base changes we count are real SNVs, because all SNVs we counted appeared as haplotypes of 3 and 5 SNVs. The extracting of reads was done with a shell script that utilizes standard NGS analysis command line packages. First, a reference sequence was generated by exporting a FASTA file from NCBI’s Genome Viewer for the mm10 mitochondrial genome set to the region of interest. This reference file was indexed for alignment using bwa index. Next, reads were aligned with bwa mem and converted to SAM format. Samtools sort and index were used to convert the reads into an organized BAM format, and samtools mpileup -uf and bcftools call -mv -0 v were used for variant calling. The reads were visualized on the IGV desktop program, where specific counts of mutants in a given position can be seen easily and with greater resolution than standard variant calling. Reads of interest were converted to a useable format for further visualization in NCBI BLAST with samtools FASTA and an in-house script for selecting a specific number of reads (as BLAST can only display up to 5,000 FASTA reads of length 133bp, so it was most efficient to portion the reads in a consistent and repeatable manner).

### Data for spatial mutational analysis (Figure 3)

Data come from the raw data files provided by the authors (e.g. NZB_Y1_Brain.dcs.vcf). These are raw counts of SNVs in ds reads without the correction for NUMTchr1.

